# Persistent hepatitis B virus and HIV coinfections in dually humanized mice engrafted with human liver and immune system

**DOI:** 10.1101/2023.05.13.540563

**Authors:** Glenn Hogan, Benjamin Y. Winer, James Ahodantin, Julie Sellau, Tiffany Huang, Florian Douam, Masaya Funaki, Luis Chiriboga, Lishan Su, Alexander Ploss

## Abstract

Chronic hepatitis B (CHB), caused by hepatitis B virus (HBV), remains a major medical problem. HBV has a high propensity for progressing to chronicity and can result in severe liver disease, including fibrosis, cirrhosis and hepatocellular carcinoma. CHB patients frequently present with viral coinfection, including HIV and hepatitis delta virus. About 10% of chronic HIV carriers are also persistently infected with HBV which can result in more exacerbated liver disease. Mechanistic studies of HBV-induced immune responses and pathogenesis, which could be significantly influenced by HIV infection, have been hampered by the scarcity of immunocompetent animal models. Here, we demonstrate that humanized mice dually engrafted with components of a human immune system and a human liver supported HBV infection, which was partially controlled by human immune cells, as evidenced by lower levels of serum viremia and HBV replication intermediates in the liver. HBV infection resulted in priming and expansion of human HLA-restricted CD8+ T cells, which acquired an activated phenotype. Notably, our dually humanized mice support persistent coinfections with HBV and HIV which opens opportunities for analyzing immune dysregulation during HBV and HIV coinfection and preclinical testing of novel immunotherapeutics.

## INTRODUCTION

Hepatitis B virus (HBV) imposes a considerable disease burden globally. Approximately 257 million people are infected chronically with HBV (CHB), and are resultantly at greater risk of developing cirrhotic liver disease and hepatocellular carcinoma ^1^. About one million deaths occur annually from CHB and, compared with 2016, projections for 2040 estimate that deaths due to liver cancer caused by HBV will almost double ^2^. Although de novo HBV infection can be effectively prevented via vaccination, and viremia stably suppressed with direct-acting antivirals (DAAs), chronic HBV is largely incurable ^3^. HBV infection often presents clinically in patients who have complex conditions and coinfections with other viruses. One such virus is the human immunodeficiency virus type 1 (HIV-1), for which there is neither a cure nor vaccine ^4^. Of the 38 million people worldwide, who are infected with HIV, approximately 10% of them are coinfected with HBV ^5^.

It is noted that HIV infection can induce immunosuppression and inflammation that may exacerbate HBV-mediated liver disease progression. However, most reports originated from studies conducted before the highly active combination anti-retroviral therapy (cART) entered widespread clinical use ^6^. Reverse transcriptase inhibitors can inhibit replication of both HIV and HBV. This has affected the way in which these viruses interface in patients receiving such treatment, and clinical studies of HBV and HIV coinfection reveal the interpretational difficulties that can arise in this setting. For example, one case study observed that lamivudine-inclusive cART had a propensity to control occult HBV infection (OBI), given that discontinuation of it resulted in OBI reactivation in a coinfected patient ^7^. Prescription of cART has been used to explain the observation that HBV coinfection in HIV-infected patients did not correlate with an increased risk of hepatotoxicity ^8^. Conversely, coinfected patients initiated on cART can experience immune reconstitution inflammatory syndrome that differs from that experienced by HBV-monoinfected patients ^9^. Some components of cART are also hepatotoxic ^10^, which further complicates the treatment of patients coinfected with HBV and HIV.

Due to a scarcity of suitable, small animal models that recapitulate human-like features of infection with these viruses, studies that explore the mechanisms that govern the pathophysiology of HBV and HIV coinfection have been confined almost exclusively to the clinic. This has resulted in a highly limited, and sometimes contradictory, understanding of how these viruses interact with each other and their host. For example, it has been noted that the immune reconstitution in response to HIV suppression by cART could enhance the rate at which seroconversion of HBV antigens occurs ^11^. It has been postulated that a cART-induced rise in CD4+ T cells could accelerate the immune response to HBV in coinfected patients ^12^, presumably by enhancing antigen presentation and thereby antibody production against HBV. Other reports question the validity of using CD4+ T-cell count as an indicator of treatment efficacy, given the evidence that patients can fail to stabilize inflammation despite significant rises in CD4+ T-cells ^13^.

Until an animal model is made that recapitulates faithfully and authentically (some aspects of) HIV-exacerbated viral hepatitis, elucidating the mechanisms underlying the common and serious diseases caused by these pathogens and the development of effective therapies will be impeded. Humanized mice, i.e. animals engrafted with human tissues and/or expressing human genes, have emerged as a versatile experimental model to study human-tropic viruses. As HBV requires a humanized liver for its life cycle, robust engraftment of human hepatocytes has been established in a number of immunodeficient liver injury models, including Alb-uPA ^14^, FAH^-/-^ ^15^, MUP-uPA ^16^, and HSV-TK ^17^ mice. The resultant human liver chimeric mice support HBV infection and have been used to study innate host responses to HBV and for testing the efficacy of novel therapeutic regimens. However, the highly immunocompromised status of these human liver chimeric mice precludes the study of immune-mediated pathogenesis by HBV.

To enable analysis of human immune responses to HBV and HIV, protocols are being designed to co-engraft mice with human hepatocytes and components of a human immune system (HIS). Double humanization of both the liver and immune system has been achieved with human hematopoietic stem cells (HSCs) and either adult ^18,19^ or fetal ^20^ hepatocytes. Maturation of fetal hepatoblasts by exogenous administration of human oncostatin M considerably boosted human hepatic chimerism, affirming that less mature hepatic cells do not proliferate in response to liver injury in these xenorecipients. Dually engrafted mice can support HBV infection, and studies have shown that viral infection triggers activation of the engrafted HIS ^19^, in particular natural killer (NK) cells ^20^ and M2 macrophages ^21^, and leads to some virally induced histopathology ^21^. A dually humanized HSV-TK mouse model has also been created for the study of liver disease induced by HIV ^22^. However, to date, a humanized mouse model has yet to be realized that can support HBV and HIV coinfection, and that maps on closely to the viral kinetics observed in patients.

As the impact of HIV coinfection on patients with CHB is ill-defined, developing a small animal model to investigate this is an urgent need. This can be accomplished by tracking HBV-specific adaptive immunity in the context of HIV coinfection. Priming and maintenance of HBV antigen-specific T-cell responses have not been clearly understood during HBV infection, which is a major impediment, considering that T cells play a critical role in immune control of HBV. Here, we demonstrate that mice co-engrafted with human liver tissue and a humanized immune system are susceptible to persistent HBV and HIV coinfection, and mount HBV-specific immune responses which can be tracked with MHC-tetramers. This new animal model holds promise for gaining mechanistic insights into virus-mediated immune dysfunction and immunopathogenesis mediated by HBV and HIV coinfection.

## RESULTS

### FNRG/A2 mice support robust dual engraftment of human hepatic and hematopoietic cells

Development of functional adaptive immune responses is limited in HIS mice by the lack of human leukocyte antigen (HLA) gene expression in mouse thymic epithelial cells. Transgenic expression of a common human MHC class I allele, HLA-A2, has been shown to significantly increase HLA-restricted human antiviral T-cell responses in Hu-HIS mice infected with the human (lympho-)tropic pathogens Epstein-Barr virus (EBV), HIV ^23^, dengue virus ^24-26^ or yellow fever virus ^27^. Thus, a transgenic FNRG mouse expressing the HLA-A*0201 allele, hereafter referred to as FNRG/A2 mice, was constructed to track HBV-specific CD8+ T cells (**Figure 1A**). To facilitate hepatic engraftment, adult FNRG/A2 mice were intrasplenically injected with adult primary human hepatocytes (PHH). Between 10-14 days thereafter, PHH-injected and non-injected FNRG/A2 mice were subjected to sublethal irradiation and injected intravenously with HLA-matched human hematopoietic stem cells (HSCs). Human hematopoietic chimerism was similar in singly and dually repopulated mice, reaching ca. 20-25% of total CD45+ leukocytes in peripheral blood by 10 weeks post HSC injection (**Figure 1B**). The human peripheral blood mononuclear cell (PBMC) fraction contained CD3+ (both CD4+ and CD8+) T cells, B cells and, to a lesser extent, NK and myeloid cells (**Supplementary Figure 1**), which is largely in line with previous reports. The frequencies of the major human leukocyte subsets – including B, CD4+ and CD8+ T cells, NK cells and various myeloid populations – in the peripheral blood, spleen and liver were similar overall (**Supplementary Figure 1**). Consistent with our previous results in FNRG mice ^28^, FNRG/A2 supported robust engraftment with PHHs as indicated by human albumin concentrations reaching >5 mg/ml (**Figure 1D**) and detection of human FAH+ cells in the liver parenchyma (**Figure 1E**), yielding an estimated human hepatic chimerism of 50-80%. Of note, there was no statistically significant difference in the human hematopoietic engraftment between singly and dually engrafted cohorts of mice. Collectively, these data are in line with reports detailing dual humanization of the hepatic and hematopoietic compartments using allogenic cell sources ^18,29^. Notably, long-term dual reconstitution, without any evidence of hepatocyte rejection by the HIS, was sustained even when the human cells were mismatched in their major histocompatibility complex ^18,29^. This latter observation is consistent with the limited HLA matching in human liver transplantations, presumably due to the tolerogenic microenvironment of the liver. This may also be due to pre-engraftment of PHHs to induce tolerance during human HSC differentiation in the dually engrafted mice.

**Figure 1.**
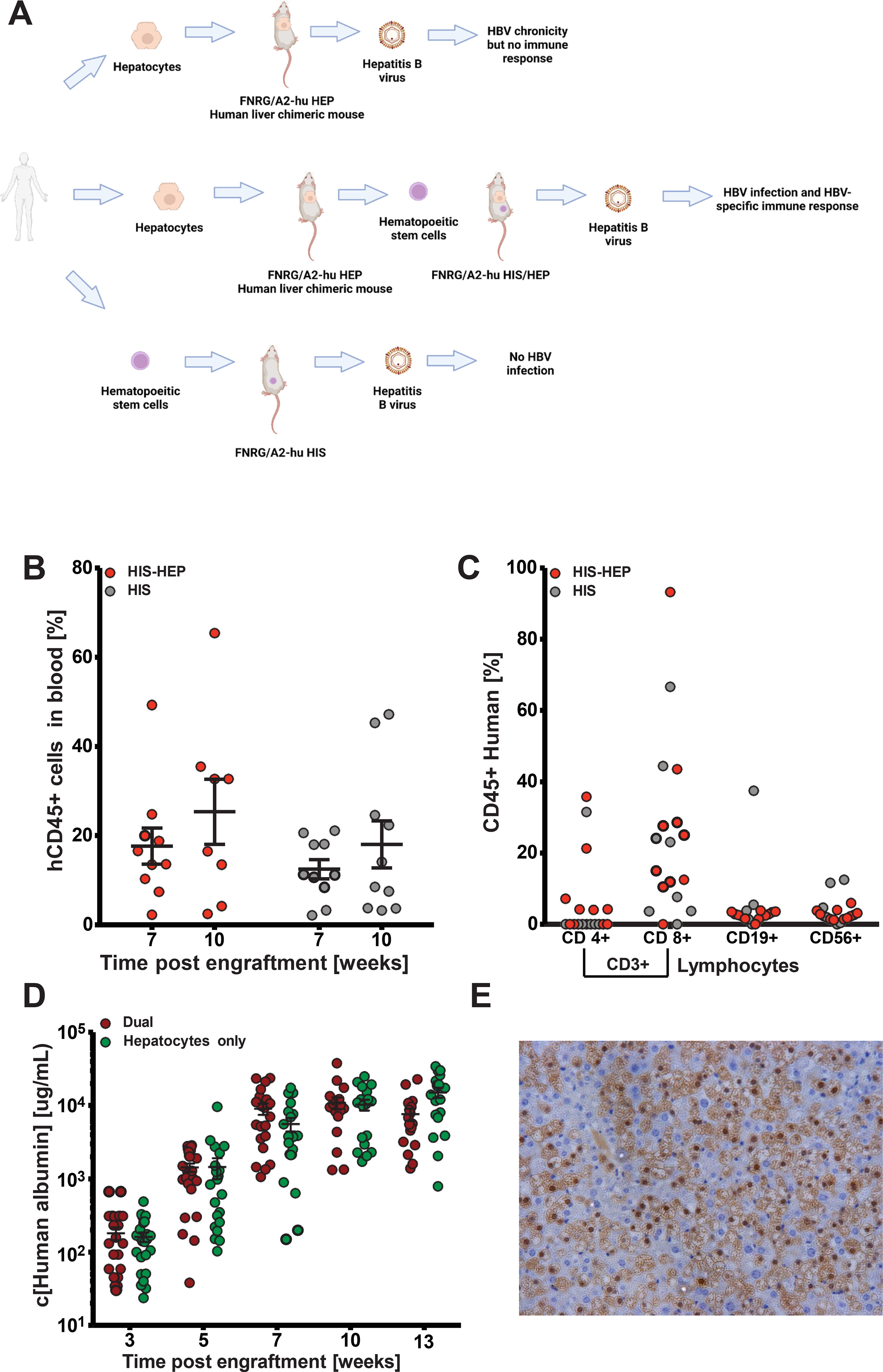
Robust human hematopoietic and hepatic co-engraftment in FNRG/A2 mice. **(A).** Schematic of overall experimental approach. FNRG/A2 mice were either dually engrafted with human HSCs and hepatocytes, or HSCs alone. (**B**). Frequency of hCD45+ cells in HIS-HEP and HIS mice. (**C**). Frequency of CD3+ T cells, CD19+ B cells, and CD56+ NK cells. HIS-HEP and Hep only mice were bled and human albumin in the serum quantified by ELISA (**D**). Human albumin concentrations in the sera of HIS-HEP vs. HEP mice. (**E**). FAH staining of HIS-HEP engrafted mice corroborates human hepatocyte engraftment. N=10 per group of HIS-HEP and HIS mice.

### Human immune cells partially control HBV infection in dually humanized mice

Next, HBV infection kinetics were characterized in hepatocyte-only (HEP), HIS-only and dually engrafted (HIS-HEP) mice. Consistent with previous reports ^14,15,28,30^, HBV viremia was readily detectable in the serum of HEP and HIS-HEP mice within 2 weeks of intravenous inoculation and plateaued at ca. 4 weeks (**Figure 2A, B**). In dually engrafted mice, HBV DNA (**Figure 2A**) and HBV surface antigen (HBsAg) (**Figure 2B**) reached similar levels by 2 weeks post infection (wpi), but viremia subsequently decreased relative to HEP-only mice, suggesting some level of control by the engrafted HIS. These data were further corroborated by significantly lower copy numbers of intrahepatic HBV DNA (**Figure 2C**) and pregenomic RNA (pgRNA) (**Figure 2D**) 6 wpi in HIS-HEP versus HEP mice. Notably, levels of covalently closed circular DNA (cccDNA), a crucial marker for HBV persistence (**Figure 2E**), were similar between the cohorts. In line with these observations, HBV core antigen (HBcAg)-expressing cells were readily detectable in the livers of HEP mice (**Figure 2F**) unlike in the HIS-HEP mice (**Figure 2G**). Expectedly, singly engrafted HIS control mice, which do not harbor HBV-permissive, human hepatocytes, remained aviremic (**Figure 2C-E**). Collectively, these data demonstrate that, in this model, an engrafted HIS can partially control HBV infection but did not clear the infection or reduce HBV cccDNA within the 6-week study period.

**Figure 2.**
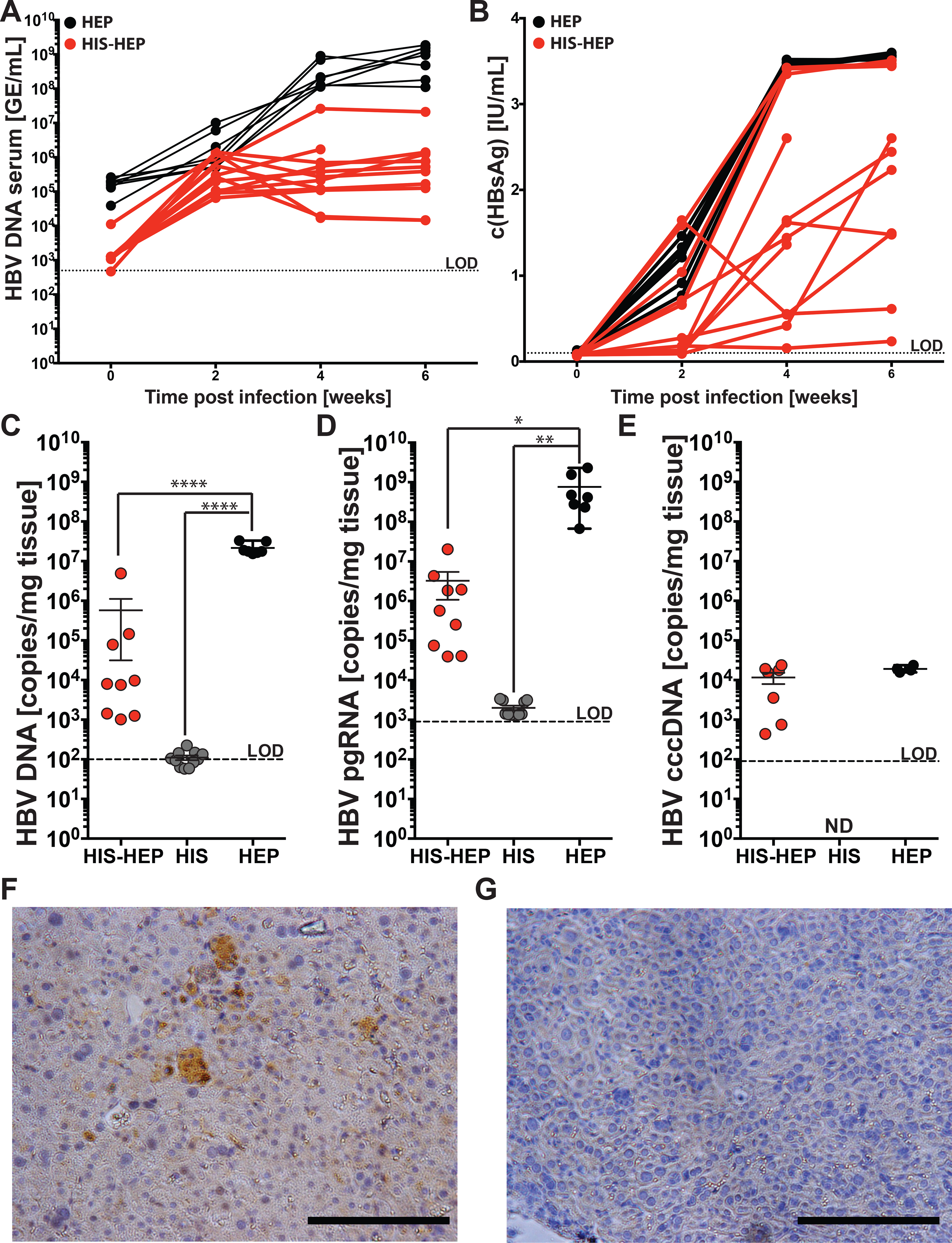
Partial control of an acute HBV infection in dually engrafted FNRG/A2-hu HIS-HEP mice. (**A/B**). HBV infection kinetics were measured in serum by HBV DNA **(A)** or HBsAg **(B)**. **(C-G).** In the liver, total HBV DNA (**C**), HBV pgRNA (**D)**, or HBV cccDNA (**E**) were quantified by qPCR. Immunohistochemical staining for HBcAg was performed on FNRG/A2-hu HIS-HEP mice (**F**) and FNRG/A2-hu HIS mice (**G**). N=7-10 animals per group. Error bars represent means ± SEM. Multiple group comparisons were analyzed by one-way ANOVA with a Bonferroni’s multiple comparisons test. *p<0.05, **p ≤ 0.01, ****p≤ 0.0001.

### HLA-restricted CD8+ T cells acquire an activated phenotype during HBV infection

To determine whether the engrafted HIS in HIS-HEP mice would respond to HBV infection and prime HBV antigen-specific T cells, cohorts of HIS and HIS-HEP mice engrafted with A2+ HSCs and A2+ PHHs were infected with HBV, and human leukocytes subjected to flow cytometric analysis. Overall, the frequencies of the major lymphoid and myeloid subsets – including B, CD4+ and CD8+ T cells, NK cells and various myeloid populations – in the peripheral blood, spleen and liver did not change significantly upon infection (**Supplementary Figure 1**). To detect and quantify the frequencies of HBV-specific T cells, an HLA-A*0201-restricted HBcAg-derived epitope (FLPSDFFPSV) tetramer was used, which can assess virus-specific T-cell immunity in HBV-infected patients ^31^. In response to HBV infection, dually engrafted HIS-HEP but not HIS mice, mounted human HLA-A*0201-restricted HBcAg-specific (FLPSDFFPSV) CD8+ T-cell responses in the spleen and livers (**Figure 3A, B**). The cell-surface phenotype of the antigen-specific cells, and thus co-stained FLPSDFFPSV:HLA-A2*0201-tetramer+ CD8+ T cells, was determined by staining for several activation and exhaustion markers, including PD1, CCR7, CD38, HLA-DR, CD45RA and CD127. Akin to data in patients, some spread in the overall levels of surface expression was observed (**Figure 3C, D**). In the liver of HIS-HEP mice, the activation markers HLA-DR1 and CD127 were significantly upregulated on tetramer+ CD8+ T cells (**Figure 3C, D**). In the spleen, only CD127 expression reached statistical significance (**Figure 3E**).

**Figure 3.**
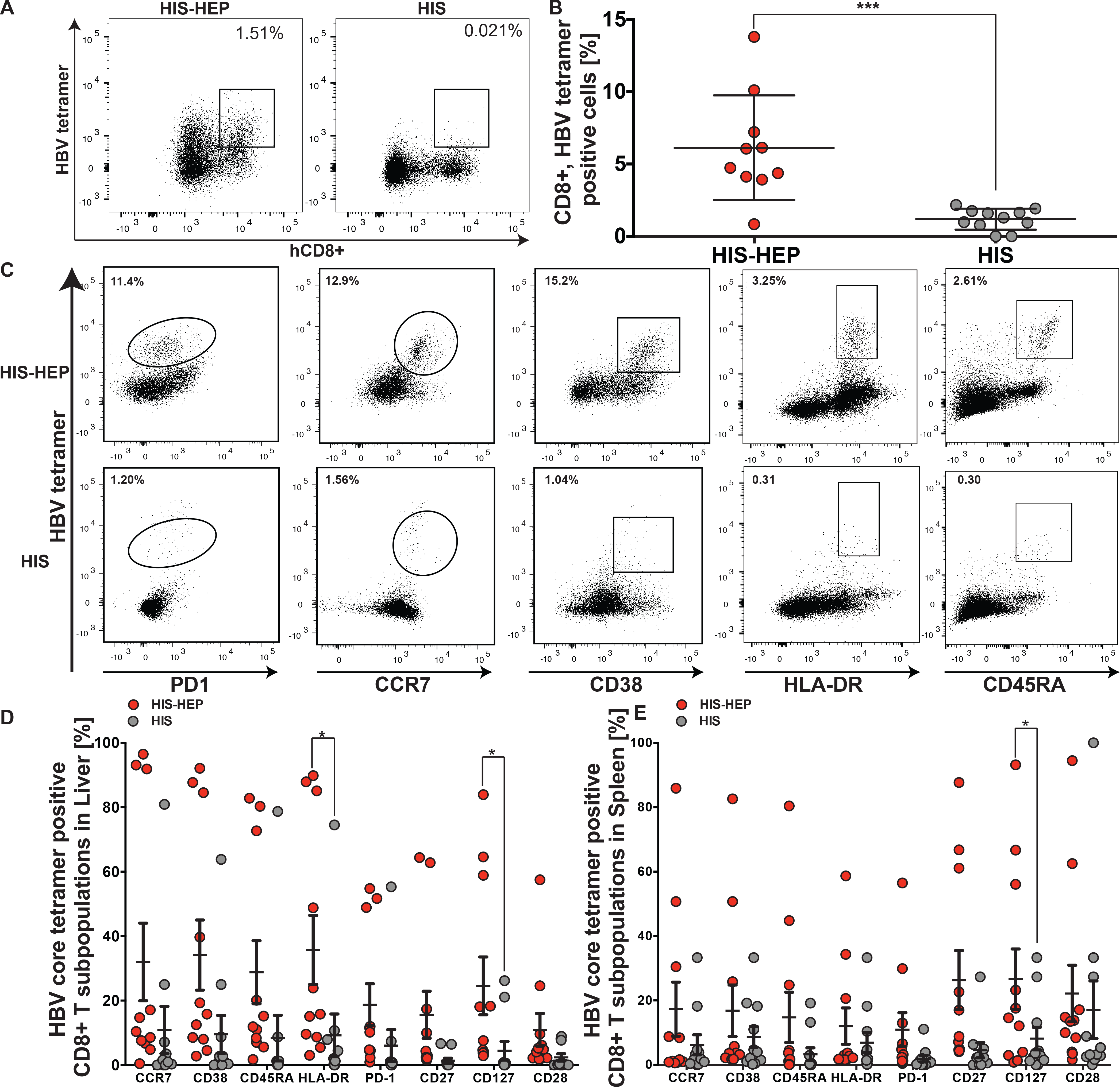
Priming and expansion of intrahepatic HBcAg-specific CD8+ T cells in HBV-infected FNRG/A2-hu HIS-HEP mice. FNRG/A2-hu HIS-HEP and FNRG/A2-hu HIS mice were challenged with HBV, and their lymphocytes were isolated from livers and spleens. (**A**). Representative FACS plots of hCD8+, HBV tetramer-positive cells. (**B**). Quantification of the number of hCD8+ T cells positive for HBV core FLPSDFFPSV/A2 tetramer staining from FNRG/A2-hu HIS-HEP (red) and FNRG/A2-hu HIS mice (gray). (**C**). Representative FACS plots of hCD8+ T cells that were dually positive for their respective activation markers (PD1, CCR7, CD38, HLA-DR, or CD45RA) and the HBV core tetramer. **(D/E).** Quantification of CD8+ T cells in the liver (**D**) and spleen (**E**) that were dually positive for respective activation marker (CCR7, CD38, CD45RA, HLA-DR, PD-1, CD27, CD127, and CD28) and the HBV tetramer. N=7-10 animals per group. Error bars represent means ± SEM. Multiple group comparisons were analyzed by one-way ANOVA with a Bonferroni’s multiple comparisons test. *p <0.05, ***p ≤ 0.001.

Collectively, these data provide evidence that HBV-specific T cells are primed during viral infection in humanized mice. Notably, previously developed dual chimeric HIS-HEP mice have only been generated using conventional immunodeficient liver injury mice and therefore were unable to develop antigen-specific responses ^18,21,29^. Thus, this novel model opens opportunities to mechanistically dissect HBV-induced immune dysfunction ^32^. In NSG/A2-hu HSC mice treated with anti-mouse Fas mAb, similar findings were reported with A2/HBc-specific CD8+ T cells. Interestingly, those tetramer+ CD8 T cells were reduced in their ability to respond to HBcAg *in vitro* relative to similar CD8+ T cells from the spleen ^21^. It will be of interest to study the functional difference between HBV-specific T cells in the spleen and in the liver.

### Limited evidence of immune-mediated liver injury during an acute HBV infection in HIS-HEP mice

Although the mechanisms underlying liver disease in HBV-infected patients are not fully understood, it is thought that the inflammatory milieu in the liver during infection is a significant driver of hepatic pathology. Thus, we sought to determine whether we could observe any evidence of HBV-induced liver injury in our model. There were no discernable differences in the overall appearance of the liver tissue and no evidence of fibrosis in either infected or non-infected HIS, HEP, or HIS-HEP mice during the acute phase of HBV infection (**Figure 4A-F**). Overt liver disease is rare in HBV-infected patients during the acute phase of the infection. The apparent lack of liver disease in our model is also largely in agreement with previous observations in dually humanized mice ^19,20^ and may be attributed to the overall impaired function of the engrafted immune system and/or the lack of human non-parenchymal cells in the liver which may aid in driving disease progression. Furthermore, liver disease is far more exacerbated in patients chronically infected with HBV for many months to years and thus the time course will likely have to be extended beyond the current 6-week study period to observe a liver injury phenotype. In NSG/A2-hu HSC mice co-transplanted with fetal hepatoblasts, and treated with anti-mouse Fas mAb, low HBV replication is detected and persistent HBV infection for 14-16 wpi leads to liver diseases associated with induction of human M2-like macrophages ^21^. It will be of interest to investigate HBV-induced liver diseases in the HIS-HEP model during chronic, persistent infection and with additional cofactors.

**Figure 4.**
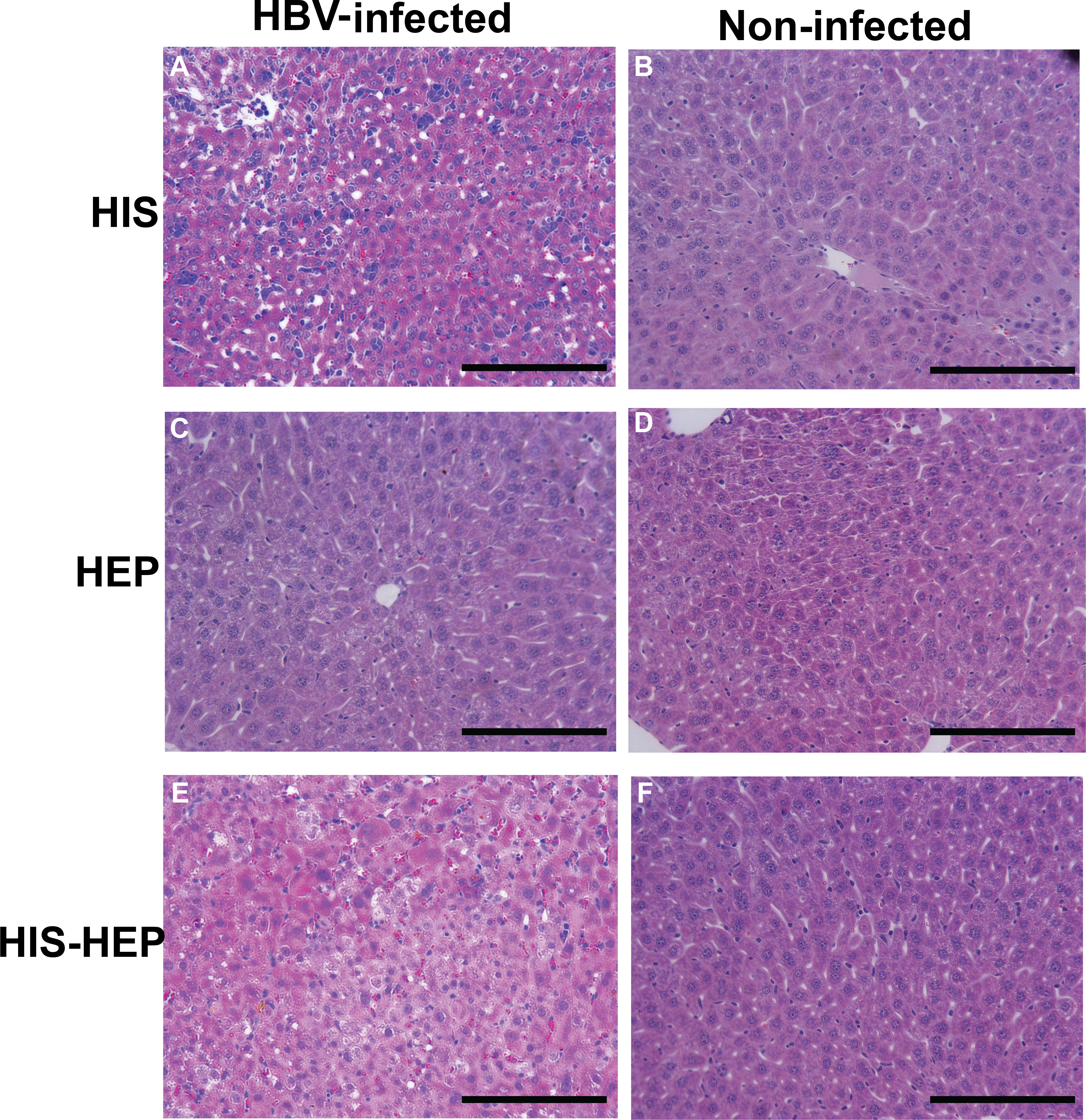
Minor liver injury is observed in HIS-HEP vs HEP mice. HBV infected HIS-HEP, HIS, and HEP mice were euthanized 6 weeks post challenge with HBV. Mouse livers were perfused and embedded in paraffin and H&E stained to ascertain if liver damage had occurred. HBV-infected HIS (**A**), HEP (**C**), and HIS-HEP (**E**) and non-infected HIS (**B**), HEP (**D**), and HIS-HEP (**F**) mice showed little evidence of liver damage.

### Dually humanized mice can support persistent coinfection with HBV and HIV-1

We aimed to determine whether dually engrafted mice would support HBV and HIV confection. It has previously been demonstrated that deletion of the murine *Flk2* gene (*Flk2*-/-) severely impairs the development of various myeloid cell types, including dendritic cells (DC) ^33^. It has also been shown by our group that exogenous administration of human Flt3LG promotes the expansion of human DCs and NK cells, while leaving such cells of murine origin unaffected in *Flk2*-/- humanized mice ^27^. Thus, FNRG/A2-hu HIS-HEP mice deficient in *Flk2* (FNRGF/A2-hu HIS-HEP) were generated. Cohorts of dually humanized FNRGF/A2-hu HIS-HEP mice were first infected intraperitoneally with HBV and subsequently with HIV. Mice were then treated with Flt3LG to promote myeloid and NK cell expansion (**Figure 5A**). These dually humanized mice supported persistent HBV and HIV coinfection, as evidenced by serum HBsAg (**Figure 5B**), HBV DNA and pgRNA in the liver (**Figure 5C**), and serum HIV RNA (**Figure 5D**). Over the six-week study period, no overt effect of HIV on HBV infection metrics was observed, and vice versa. However, differences may become more apparent over longer periods of coinfection, and inclusion of additional parameters (i.e. functionality of T cells, number of HIV+ cells, mutation rate in either virus, etc.). Collectively, this FNRGF/A2-hu HIS-HEP mouse model is well suited for such studies.

**Figure 5.**
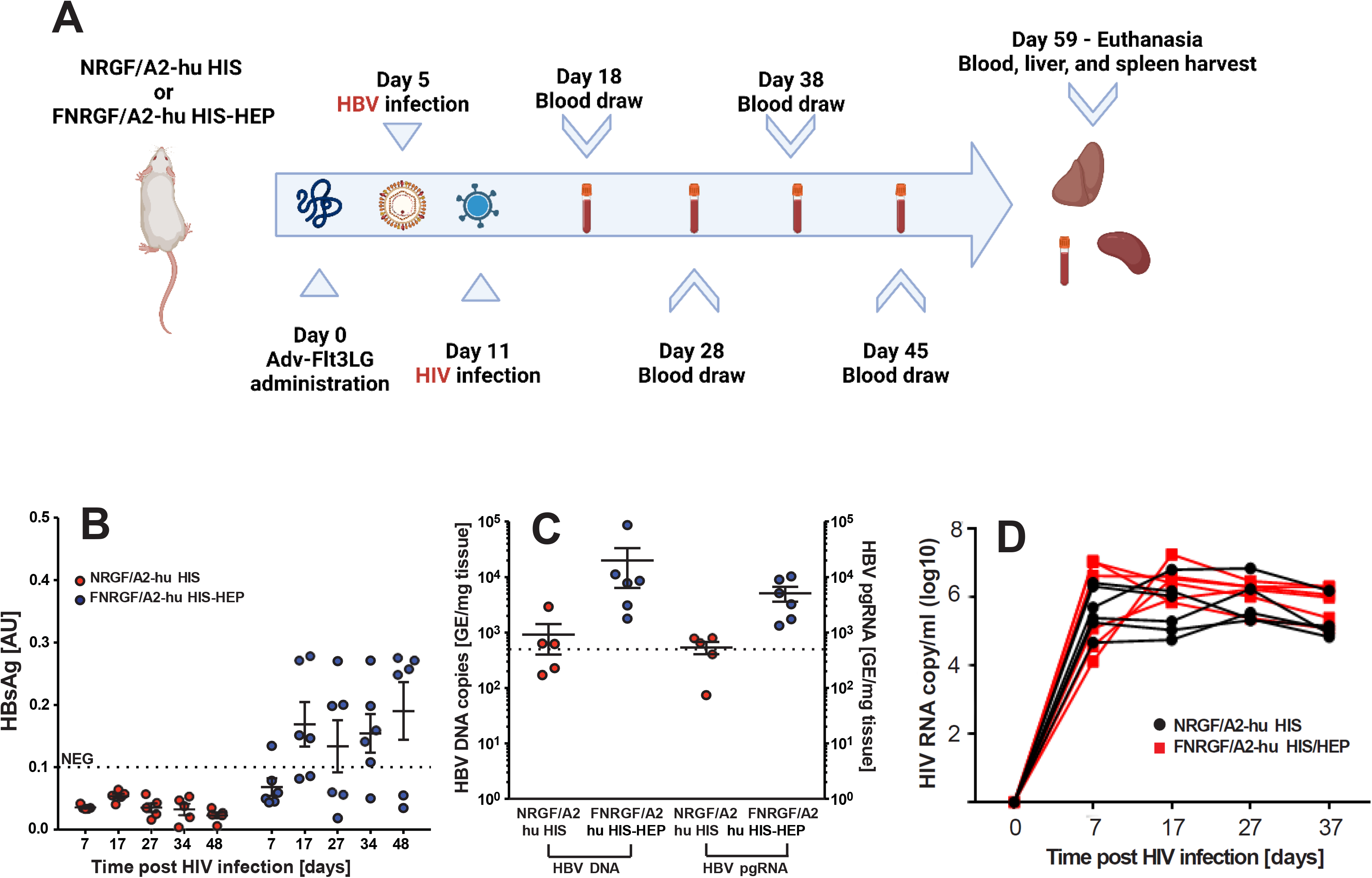
HBV/HIV-1 coinfection in FNRGF/A2-hu HIS-HEP mice. **(A.)** Schematic representation of the experimental procedure for characterizing HBV and HIV-1 infection in NRGF/A2-hu HIS mice and FNRGF/A2-hu HIS-HEP mice. (**B.)** Longitudinal HBsAg in the serum of NRGF/A2-hu HIS (red) and FNRGF/A2-hu HIS-HEP (blue) mice following HBV and HIV coinfection. (**C.)** HBV DNA and pgRNA copies per mg liver tissue in coinfected NRGF/A2-hu HIS (red) and FNRGF/A2-hu HIS-HEP (blue) at termination (48 days post-HIV-1 infection and 54 days post-HBV infection). (**D.)** HIV viral load in the serum of coinfected NRGF/A2-hu HIS (red) and FNRGF/A2-hu HIS-HEP mice (black) during the first 37/31 days post HBV/HIV-1 coinfection, respectively. **p<0.01, ****p<0.0001.

### Dually humanized mice experience coinfection-dependent differences in viral titer and humanized immunity

Finally, we aimed to determine if our model can recapitulate fundamental features of HBV and HIV coinfection that are observed clinically. To do this, we examined our FNRGF/A2-hu HIS-HEP model under various settings of infection, including uninfected mice and those singly and coinfected with HBV and/or HIV. Mice to be coinfected were infected first with HBV, followed by HIV, and all mice were bled several times over 15 weeks, followed by liver harvest **(Figure 6A)**. The presence of HBV DNA in the blood of coinfected mice and those singly infected with HBV **(Figure 6B)**, and the presence HIV RNA in coinfected mice and those singly infected with HIV **(Figure 6C)** was then confirmed. HIV and HBV did not significantly affect each other’s replication, as determined by viremias in the blood.

**Figure 6.**
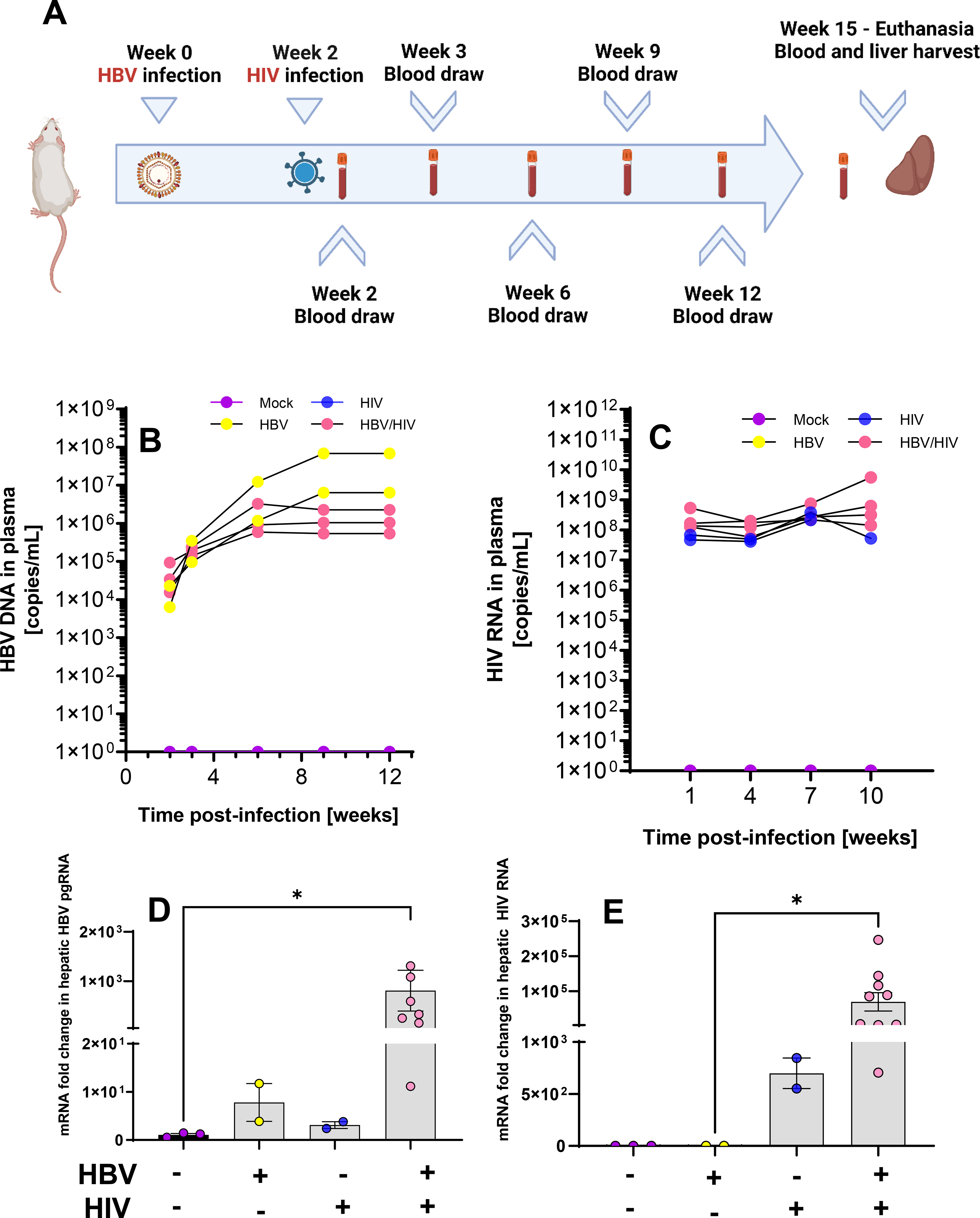
HBV and HIV infection and replication in FNRGF/A2-hu HIS-HEP mice infected with HBV and/or HIV. **(A.)** Schematic representation of the experimental procedure for single infection with HBV or HIV, and coinfection. (**B.)** Quantification of HBV genomic DNA in the plasma of singly infected, coinfected, and uninfected mice. (**C.)** Quantification of HIV genomic RNA in the plasma of singly infected, coinfected, and uninfected mice. (**D.)** Quantification of HBV pgRNA in the livers of singly infected, coinfected, and uninfected mice (relative to human GAPDH). (**E.)** Quantification of HIV RNA in the livers of singly infected, coinfected, and uninfected mice (relative to human GAPDH). Error bars represent means ± SEM. Significance was determined via the Kruskal-Wallis test. *p<0.05.

We next quantified changes in gene expression in terms of HBV pgRNA **(Figure 6D)** and HIV RNA **(Figure 6E)** in the livers of these animals. While no significant difference was calculated in terms of HBV pgRNA expression in the livers of HBV-monoinfected mice compared with coinfected mice, changes in expression ranging from hundred-fold to thousand-fold were frequently recorded in the liver of coinfected animals. This substantiates what is known clinically about the exacerbation of HBV-related disease in coinfected patients not receiving cART. Similarly, no significant difference was detected in terms of HIV RNA expression in the livers of HIV-monoinfected mice compared with coinfected mice; however, a trend in increased HIV gene expression/replication in coinfected mice was also noted.

HBV/HIV coinfections are known to result in more severe liver disease than HBV mono-infections, and thus we aimed to gather evidence of whether such a phenotype could possibly be recapitulated in our dually humanized mice. We thus quantified the expression of various human genes, related to inflammation and liver disease, by RT-qPCR in the livers of these animals (**Figure 7**). This analysis included human macrophage markers and fibrosis genes – CD163, arginase 1 (ARG1), tissue inhibitor of metalloproteinase-1 (TIMP-1), and transforming growth factor beta (TGF-β1), in addition to several interferon (IFN)-stimulated genes (ISGs), including interferon-induced transmembrane protein 3 (IFITM3), ISG15, IFN-β, myxovirus resistance protein 1 (Mx1), 2’-5’-oligoadenylate synthetase 1 (OAS1), and the cytidine deaminase APOBEC3G (APO3G). These markers are commonly analyzed in the contexts of HIV infection and coinfection with HBV ^34,35^. HBV is considered a stealth virus that does not result in significant dysregulation of host inflammation in the liver ^36^, which is also corroborated by our preliminary data (**Figure 7)**. Remarkably, there was a trend towards more pronounced expression of all 10 genes in HBV/HIV coinfected mice, as compared with the naïve and singly infected mice, but only hOAS1 expression reached statistical significance between the HBV and HBV/HIV coinfection groups (**Figure 7C**). Collectively, our data suggest that HBV/HIV co-infection results in dysregulation of numerous host genes in the liver of our dually humanized mice that mirrors what is observed in coinfected patients.

**Figure 7.**
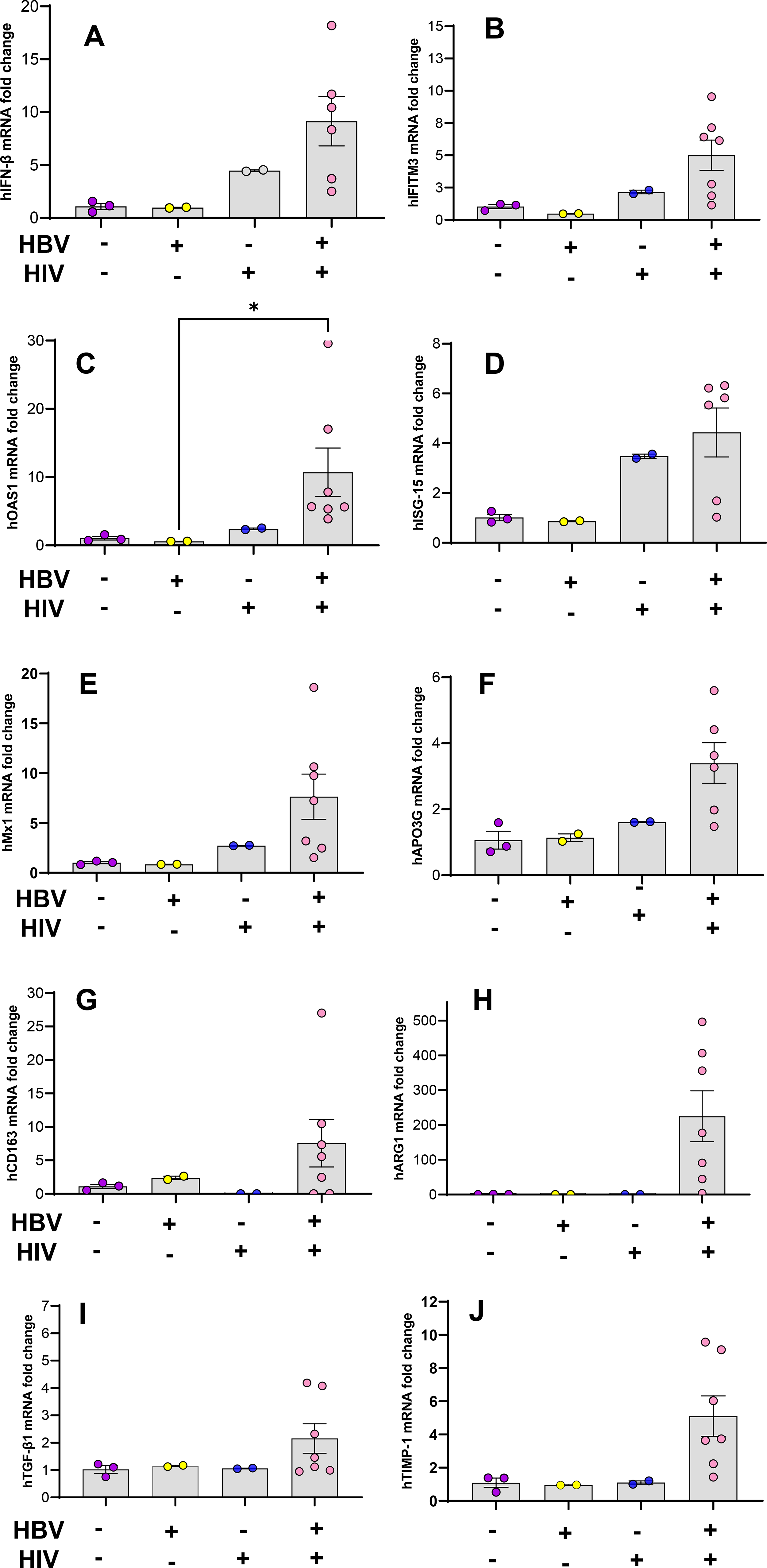
Induction of human interferon-stimulated genes (ISG), and human M2 macrophage and fibrosis genes in the livers of FNRGF/A2-hu HIS-HEP mice singly infected or coinfected with HBV and/or HIV, and uninfected mice. **(A-F).** ISGs investigated were **(A)** IFN-β, **(B)** IFITM3, **(C)** OAS1, **(D)** ISG-15, **(E)** Mx1, and **(F)** APO3G. **(G-J).** Macrophage and fibrosis genes investigated were **(G)** CD163, **(H)** ARG1, **(I)** TGF-β1, and **(J)** TIMP-1. Relative gene expression is relative to human GAPDH. Error bars represent means ± SEM. Significance was determined via the Kruskal-Wallis test. *p<0.05.

## DISCUSSION

The complex interplay between HBV, HIV, and the human host remains opaque in part due to the limited availability of immunocompetent animal models. Some basic and central questions are controversial, such as whether HBV can influence HIV pathogenesis at all. Some studies note that HIV is apparently unaffected by HBV coinfection ^37^, while others have observed that patients positive for HBeAg experience slower responses to cART ^38^. Furthermore, *in vitro* studies have shown that the X-protein of HBV can super-induce HIV-1 replication ^39^, which corroborates what is observed in our model. The lack of fundamental understanding of the true nature of HBV/HIV coinfection calls urgently for a reproducible, robust model that can accurately map onto the clinical reality. The dually engrafted models described here hold promise for addressing elusive and fundamental questions in HBV and HIV coinfection virology and pathogenesis.

One of the ways in which the impact of HIV on HBV infection might be assessed is by tracking T cells specific for HBV antigens. Immune tolerance and development of a chronic HBV infection are thought to result, at least in part, from a dysfunctional CD8+ T-cell response ^40^. In chimpanzees experimentally infected with HBV, it was shown that an HBV-specific CD8+ T-cell response was key to viral clearance ^41^. It has also been observed that CD8+ T cells from chronically infected patients are more prone to having an exhausted phenotype as indicated by high levels of programmed death 1 (PD1) molecule, while levels of PD1 are low in acutely infected patients ^42^. In other reports, CD8+ T cells in chronically infected patients have been shown to express higher levels of TIM3 and cytotoxic T-lymphocyte antigen-4 (CTLA-4) than those of acutely infected or naïve patients, also indicating an exhausted phenotype ^43^. In addition, a lower frequency of HBV-specific tetramer-stained CD8+ T cells have been isolated from chronic as opposed to acutely infected patients ^42^. Notably, *ex vivo* CD8+ T cells from chronically infected patients or HBV transgenic mice exhibit a less activated phenotype and are impaired in the ability to produce effector cytokines ^42,44^. The critical importance of the CD8+ T-cell response has been further confirmed as CD8+ T cells, engineered to recognize HBV antigens when adoptively transferred into human liver chimeric uPA/SCID mice, resulted in reduced viremia in both the serum and liver, indicating the ability of these engineered CD8+ T cells to control HBV infection ^45^. As an HBV-specific CD8+ T-cell response in HBV infected dually engrafted FNRG/A2 animals was observed in the present work, further characterization of this phenotype and the kinetics of this response in our HBV/HIV coinfection animal model could shed light on the parameters affecting CD8+ T cell functions in coinfected patients.

Undoubtedly, humanized mice cannot (yet) perfectly mimic the complex situation of chronic HBV and HIV coinfection in patients, but important aspects of it can be recapitulated, as shown here. Future refinements will focus on improving the limited functionality of the engrafted HIS. Numerous strategies to do this have been proposed (reviewed in ^46^) as several human cell lineages remain underrepresented in part due to the orthologs of non-redundant cytokines, which exhibit limited biological cross-reactivity. Our lab has previously shown that selective expansion of under-represented cell types, such as dendritic cells, NK cells and granulocytes, leads to markedly improved immune responses to the yellow fever virus (YFV) vaccine akin to those observed in YFV vaccines ^27^. Additionally, the development of functional adaptive immune responses is limited by the lack of HLA gene expression. Expressing a human MHC class I allele has multiple benefits as it allows for more faithful development of CD8+ T cells in the thymus, enables recognition of (viral) antigens in peripheral tissues by human CD8+ T cells and facilitates tracking of antigen-specific CD8+ T cells with MHC multimers, as previously shown for EBV, dengue virus ^24-26^ and, here in our study for the first time, HBV.

A shortcoming of our animal models is the lack of human MHC class II expression, which may result in CD4+ T-cell lineage disfunctions. It has previously been suggested that expression of a human MHC class II molecule, HLA-DR4, partially improves the development of functional human T and B cells ^47^. Our lab previously characterized adaptive immune responses to adenovirus infections in humanized HLA-A*0201 and HLA-DRB*01 doubly transgenic mice, finding statistically significant clearance of viral antigens from the liver ^48^. Thus, combining the FNRGF/A2 mice with a human MHC II transgenic model may be an immediate improvement. This would be particularly important when investigating the impact of HIV on CD4+ T cell exhaustion, and their function in helping B cells to induce HBsAg antibodies during HBV functional cure. As such, the immune response in HBV/HIV-challenged, dually engrafted, human MHC class II-expressing FNRGF/A2 mice could more faithfully recapitulate the response observed in human patients. Co-engraftment of improved xenorecipient strains with additional HSC donor-matched human tissues, such as liver, thymus and/or lymph nodes, could also significantly augment the immune response. Such co-engraftments could enhance T- and B-cell selection, intra-hepatic T-cell priming ^49^ and liver-mediated secretion of key human-immune components ^50^. Finally, engraftment of second-generation humanized mice with a human-like microbiome represents another valuable approach to enhance immunity, as recently suggested ^51^.

While the above-mentioned modifications will undoubtedly aid in improving our FNRGF/A2 dually humanized mouse model, the current FNRGF/A2-hu HIS-HEP mouse has already considerable utility. Here, we show that these dually engrafted animals can elicit an HBV-specific T-cell response phenotypically similar to what is observed in acutely infected HBV patients, and can sustain long-term, clinically relevant HBV and HIV coinfection. We have demonstrated that our humanized mouse model can be used effectively to interrogate transcriptional changes that occur between mono-infection and coinfection with HBV and HIV, that could influence hepatic disease progression in patients coinfected with these viruses. By cross-examining changes in liver histology with dysregulated expression of these genes, for example, we can gain keen insights into how HBV and HIV cooperate to elicit different mechanisms of disease than when they singly infect patients. Although we did not see any significant signs of liver damage or disease, this is most likely because the current study was limited to a relatively short period of time whereas it takes months to years for chronic HBV infection to result in liver disease in humans. Of note, previous work in a different type of dually engrafted mice observed that inflammatory M2 macrophages were more abundant in mice dually engrafted with human hepatocytes and a HIS as compared with non-infected dually engrafted mice ^21^. Future studies in our model or further refined versions will probe whether HBV/HIV coinfection results in accelerated liver disease mediated by the aforementioned M2-like macrophages or other lymphoid cell populations.

## MATERIALS AND METHODS

A detailed description of the Materials and Methods used in this study is included in the supplementary information.

## Authors’ contributions

G.H., B.Y.W., J.A., L.S., and A.P. designed and performed experiments and wrote the manuscript. T.S.H., F.D., M.F., and J.S. performed experiments and analysis. L.C. performed experiments.

## Acknowledgments

HepG2.2.15 cells were kindly provided by Christoph Seeger (Fox Chase Cancer Center, FCCC). We thank Gabriela Hrebikova for technical assistance and Christina DeCoste and the Molecular Biology Flow Cytometry Resource Facility, which is partially supported by the Cancer Institute of New Jersey Cancer Center Support Grant (P30CA072720. We are grateful to Jenna Gaska, Florian Douam, and members of the Ploss lab for critical discussions and edits of this manuscript. This study was supported by grants from the National Institutes of Health (R01 AI138797 to A.P. and L.S., R01 AI153236, R01 AI146917, R01 AI168048 all to A.P), a Research Scholar Award from the American Cancer Society (RSG-15-048-01-MPC to A.P.), a Burroughs Wellcome Fund Award for Investigators in Pathogenesis (to A.P.) a Graduate fellowship from the Health Grand Challenge from the Global Health Fund of Princeton University (to B.Y.W.). The NYU Experimental Pathology Immunohistochemistry Core Laboratory is supported in part by the Laura and Isaac Perlmutter Cancer Center Support Grant; NIH /NCI P30CA016087” and the National Institutes of Health S10 Grants; NIH/ORIP S10OD01058 and S10OD018338. B.Y.W. was a recipient of F31 NIH/NRSA Ruth L. Kirschstein Predoctoral awarded from the NIAID and a graduate fellowship from the New Jersey Commission on Cancer Research and was supported by an NIH training grant (T32GM007388). J.S. was a recipient of a postdoctoral fellowships from the German Research Foundation.

## REFERENCES

1. Asselah T, Loureiro D, Boyer N, Mansouri A. Targets and future direct-acting antiviral approaches to achieve hepatitis B virus cure. Lancet Gastroenterol Hepatol. 2019;4(11):883–892.

2. Foreman KJ, Marquez N, Dolgert A, et al. Forecasting life expectancy, years of life lost, and all-cause and cause-specific mortality for 250 causes of death: reference and alternative scenarios for 2016-40 for 195 countries and territories. Lancet. 2018;392(10159):2052–2090.

3. Martinez MG, Villeret F, Testoni B, Zoulim F. Can we cure hepatitis B virus with novel direct-acting antivirals? Liver Int. 2020;40 Suppl 1:27–34.

4. Davenport MP, Khoury DS, Cromer D, Lewin SR, Kelleher AD, Kent SJ. Functional cure of HIV: the scale of the challenge. Nat Rev Immunol. 2019;19(1):45–54.

5. Platt L, French CE, McGowan CR, et al. Prevalence and burden of HBV co-infection among people living with HIV: A global systematic review and meta-analysis. J Viral Hepat. 2020;27(3):294–315.

6. Kim HN. Chronic Hepatitis B and HIV Coinfection: A Continuing Challenge in the Era of Antiretroviral Therapy. Curr Hepatol Rep. 2020;19(4):345–353.

7. Costantini A, Marinelli K, Biagioni G, et al. Molecular analysis of hepatitis B virus (HBV) in an HIV co-infected patient with reactivation of occult HBV infection following discontinuation of lamivudine-including antiretroviral therapy. BMC Infect Dis. 2011;11:310.

8. Núñez M, Lana R, Mendoza JL, Martín-Carbonero L, Soriano V. Risk factors for severe hepatic injury after introduction of highly active antiretroviral therapy. J Acquir Immune Defic Syndr. 2001;27(5):426–431.

9. Mitsumoto F, Murata M, Kato Y, et al. Hepatitis B virus-related immune reconstitution inflammatory syndrome in two patients coinfected with human immunodeficiency virus diagnosed with a liver biopsy. Intern Med. 2014;53(18):2165–2170.

10. Sulkowski MS. Viral hepatitis and HIV coinfection. J Hepatol. 2008;48(2):353–367.

11. Miailhes P, Trabaud MA, Pradat P, et al. Impact of highly active antiretroviral therapy (HAART) on the natural history of hepatitis B virus (HBV) and HIV coinfection: relationship between prolonged efficacy of HAART and HBV surface and early antigen seroconversion. Clin Infect Dis. 2007;45(5):624–632.

12. Jiang T, Su B, Song T, et al. Immunological Efficacy of Tenofovir Disproxil Fumarate-Containing Regimens in Patients With HIV-HBV Coinfection: A Systematic Review and Meta-Analysis. Front Pharmacol. 2019;10:1023.

13. Tincati C, Mondatore D, Bai F, d’Arminio Monforte A, Marchetti G. Do Combination Antiretroviral Therapy Regimens for HIV Infection Feature Diverse T-Cell Phenotypes and Inflammatory Profiles? Open Forum Infect Dis. 2020;7(9):ofaa340.

14. Meuleman P, Libbrecht L, De Vos R, et al. Morphological and biochemical characterization of a human liver in a uPA-SCID mouse chimera. Hepatology. 2005;41(4):847–856.

15. Bissig KD, Wieland SF, Tran P, et al. Human liver chimeric mice provide a model for hepatitis B and C virus infection and treatment. J Clin Invest. 2010;120(3):924–930.

16. Tesfaye A, Stift J, Maric D, Cui Q, Dienes HP, Feinstone SM. Chimeric mouse model for the infection of hepatitis B and C viruses. PLoS One. 2013;8(10):e77298.

17. Kosaka K, Hiraga N, Imamura M, et al. A novel TK-NOG based humanized mouse model for the study of HBV and HCV infections. Biochem Biophys Res Commun. 2013;441(1):230–235.

18. Gutti TL, Knibbe JS, Makarov E, et al. Human hepatocytes and hematolymphoid dual reconstitution in treosulfan-conditioned uPA-NOG mice. The American journal of pathology. 2014;184(1):101–109.

19. Dusseaux M, Masse-Ranson G, Darche S, et al. Viral Load Affects the Immune Response to HBV in Mice With Humanized Immune System and Liver. Gastroenterology. 2017;153(6):1647–1661 e1649.

20. Billerbeck E, Mommersteeg MC, Shlomai A, et al. Humanized mice efficiently engrafted with fetal hepatoblasts and syngeneic immune cells develop human monocytes and NK cells. J Hepatol. 2016;65(2):334–343.

21. Bility MT, Cheng L, Zhang Z, et al. Hepatitis B virus infection and immunopathogenesis in a humanized mouse model: induction of human-specific liver fibrosis and M2-like macrophages. PLoS Pathog. 2014;10(3):e1004032.

22. Dagur RS, Wang W, Makarov E, Sun Y, Poluektova LY. Establishment of the Dual Humanized TK-NOG Mouse Model for HIV-associated Liver Pathogenesis. J Vis Exp. 2019(151).

23. Cheng L, Ma J, Li G, Su L. Humanized Mice Engrafted With Human HSC Only or HSC and Thymus Support Comparable HIV-1 Replication, Immunopathology, and Responses to ART and Immune Therapy. Front Immunol. 2018;9:817.

24. Shultz LD, Saito Y, Najima Y, et al. Generation of functional human T-cell subsets with HLA-restricted immune responses in HLA class I expressing NOD/SCID/IL2r gamma(null) humanized mice. Proc Natl Acad Sci U S A. 2010;107(29):13022–13027.

25. Strowig T, Gurer C, Ploss A, et al. Priming of protective T cell responses against virus-induced tumors in mice with human immune system components. J Exp Med. 2009;206(6):1423–1434.

26. Jaiswal S, Pearson T, Friberg H, et al. Dengue virus infection and virus-specific HLA-A2 restricted immune responses in humanized NOD-scid IL2rgammanull mice. PLoS One. 2009;4(10):e7251.

27. Douam F, Ziegler CGK, Hrebikova G, et al. Selective expansion of myeloid and NK cells in humanized mice yields human-like vaccine responses. Nat Commun. 2018;9(1):5031.

28. Winer BY, Huang T, Low BE, et al. Recapitulation of treatment response patterns in a novel humanized mouse model for chronic hepatitis B virus infection. Virology. 2017;502:63–72.

29. Wilson EM, Bial J, Tarlow B, et al. Extensive double humanization of both liver and hematopoiesis in FRGN mice. Stem Cell Res. 2014;13(3 Pt A):404-412.

30. Dandri M, Burda MR, Torok E, et al. Repopulation of mouse liver with human hepatocytes and in vivo infection with hepatitis B virus. Hepatology. 2001;33(4):981–988.

31. Bengsch B, Martin B, Thimme R. Restoration of HBV-specific CD8+ T cell function by PD-1 blockade in inactive carrier patients is linked to T cell differentiation. J Hepatol. 2014;61(6):1212–1219.

32. Tran TT. Immune tolerant hepatitis B: a clinical dilemma. Gastroenterol Hepatol (N Y). 2011;7(8):511–516.

33. Waskow C, Liu K, Darrasse-Jèze G, et al. The receptor tyrosine kinase Flt3 is required for dendritic cell development in peripheral lymphoid tissues. Nat Immunol. 2008;9(6):676–683.

34. Ahodantin J, Nio K, Funaki M, et al. Type I interferons and TGF-β cooperate to induce liver fibrosis during HIV-1 infection under antiretroviral therapy. JCI Insight. 2022;7(13).

35. Cheng L, Yu H, Li G, et al. Type I interferons suppress viral replication but contribute to T cell depletion and dysfunction during chronic HIV-1 infection. JCI Insight. 2017;2(12).

36. Winer BY, Gaska JM, Lipkowitz G, et al. Analysis of Host Responses to Hepatitis B and Delta Viral Infections in a Micro-scalable Hepatic Co-culture System. Hepatology. 2020;71(1):14–30.

37. Chun HM, Mesner O, Thio CL, et al. HIV outcomes in Hepatitis B virus coinfected individuals on HAART. J Acquir Immune Defic Syndr. 2014;66(2):197–205.

38. Idoko J, Meloni S, Muazu M, et al. Impact of hepatitis B virus infection on human immunodeficiency virus response to antiretroviral therapy in Nigeria. Clin Infect Dis. 2009;49(8):1268–1273.

39. Gómez-Gonzalo M, Carretero M, Rullas J, et al. The hepatitis B virus X protein induces HIV-1 replication and transcription in synergy with T-cell activation signals: functional roles of NF-kappaB/NF-AT and SP1-binding sites in the HIV-1 long terminal repeat promoter. J Biol Chem. 2001;276(38):35435–35443.

40. Ye B, Liu X, Li X, Kong H, Tian L, Chen Y. T-cell exhaustion in chronic hepatitis B infection: current knowledge and clinical significance. Cell Death Dis. 2015;6:e1694.

41. Thimme R, Wieland S, Steiger C, et al. CD8(+) T cells mediate viral clearance and disease pathogenesis during acute hepatitis B virus infection. J Virol. 2003;77(1):68–76.

42. Boni C, Fisicaro P, Valdatta C, et al. Characterization of hepatitis B virus (HBV)-specific T-cell dysfunction in chronic HBV infection. J Virol. 2007;81(8):4215–4225.

43. Schurich A, Khanna P, Lopes AR, et al. Role of the coinhibitory receptor cytotoxic T lymphocyte antigen-4 on apoptosis-Prone CD8 T cells in persistent hepatitis B virus infection. Hepatology. 2011;53(5):1494–1503.

44. Wang Q, Pan W, Liu Y, et al. Hepatitis B Virus-Specific CD8+ T Cells Maintain Functional Exhaustion after Antigen Reexposure in an Acute Activation Immune Environment. Front Immunol. 2018;9:219.

45. Kah J, Koh S, Volz T, et al. Lymphocytes transiently expressing virus-specific T cell receptors reduce hepatitis B virus infection. J Clin Invest. 2017;127(8):3177–3188.

46. Walsh NC, Kenney LL, Jangalwe S, et al. Humanized Mouse Models of Clinical Disease. Annu Rev Pathol. 2017;12:187–215.

47. Suzuki M, Takahashi T, Katano I, et al. Induction of human humoral immune responses in a novel HLA-DR-expressing transgenic NOD/Shi-scid/gammacnull mouse. Int Immunol. 2012;24(4):243–252.

48. Billerbeck E, Horwitz JA, Labitt RN, et al. Characterization of human antiviral adaptive immune responses during hepatotropic virus infection in HLA-transgenic human immune system mice. J Immunol. 2013;191(4):1753–1764.

49. Huang LR, Wohlleber D, Reisinger F, et al. Intrahepatic myeloid-cell aggregates enable local proliferation of CD8(+) T cells and successful immunotherapy against chronic viral liver infection. Nature immunology. 2013;14(6):574–583.

50. Sander LE, Sackett SD, Dierssen U, et al. Hepatic acute-phase proteins control innate immune responses during infection by promoting myeloid-derived suppressor cell function. J Exp Med. 2010;207(7):1453–1464.

51. Gulden E, Vudattu NK, Deng S, et al. Microbiota control immune regulation in humanized mice. JCI insight. 2017;2(21).

