## supplementary information for "Persistent hepatitis B virus and HIV coinfections in dually humanized mice engrafted with human liver and immune system"

for

**Supplementary Figure 1. Many myeloid lymphocyte lineages arise from human HSC engraftment in FNRG-A2 HIS-HEP and HIS mice.** FNRG-A2-HIS-HEP and -HIS mice were euthanized, lymphocytes isolated from their blood, spleen, and liver and stained with various immune markers. Shown are frequencies of the indicated immune cell populations. N=7-10 animals per group. Error bars represent means ± SEM. Multiple group comparisons were analyzed by one-way ANOVA with a Bonferroni's multiple comparisons test. p values *= <0.05, ** ≤ 0.01.

**MATERIALS AND METHODS**

**Mice**

The generation of *Fah*-/- NOD.Cg-*Rag1^tm1Mom^IL2rg^tmlWjl^/*SzJ IL2Rg^null^ (FNRG) mice has been previously described ^1^. FNRG expressing a human HLA-A*0201 transgene (FNRG/A2) were generated by intercrossing FNRG with NRG-Tg (HLA-A2.1)1Eng/Sz) mice ^2^ and genotyping offspring with primers that distinguish wild-type and mutant alleles. To discourage expansion of murine myeloid cell lineages, a null allele was introduced into NRG/A2 mice for the gene encoding the receptor Fms-like tyrosine kinase 3 (Flt3 or Flt2), yielding NRGF mice ^3^. FNRGF/A2 mice were produced by backcrossing FNRG mice with NRGF/A2 mice. All animal experiments were performed in accordance with a protocol reviewed and approved by the Institutional Animal Care and Use Committee (IACUC) of Princeton University (protocol number 3063).

**Human hepatocyte engraftment**

FNRG and FNRG/A2 mice were generated and transplanted as previously described ^4^ ^1^. Female mice greater than 6 weeks of age were transplanted with ca. 1 x 10^6^ cryopreserved adult human hepatocytes. Primary hepatocytes were obtained from Bioreclamation (Westbury, NY). FNRG mice were cycled on NTBC (Yecuris Inc., Tualatin, OR) supplemented in their water to block the accumulation of toxic metabolites. All surgical experiments were performed in accordance with a protocol reviewed and approved by the IACUC of Princeton University (protocol number 1930-16).

**Isolation of human CD34+ hematopoietic stem cells**

Human fetal livers (16-22 weeks of gestational age) were procured from Advanced Bioscience Resources (ABR), Inc. (Alameda, CA). Fetal liver was homogenized and incubated in digestion medium (HBSS with 0.1% collagenase IV (Sigma- Aldrich, Darmstadt, Germany), 40-mM HEPES, 2-M CaCl_2_ and 2-U/ml DNase I (Roche, Basel, Switzerland) for 30 min at 37°C. Human CD34+ HSCs were isolated using a CD34+ HSC isolation kit (Stem Cell Technologies, Vancouver, British Columbia, Canada), according to the manufacturer’s protocol. Purification of human CD34+ cells was assessed by quantification via flow cytometry, using an anti-human CD34+-FITC antibody (dilution 1:100, clone 581, BD Biosciences, Franklin Lakes, NJ). Expression of human CD90, CD38, CD45RA, and HLA-A*0201 was assessed among the CD34+ population. All experiments were performed with authorization from the Institutional Review Board and the IACUC of Princeton University.

**Human hematopoietic stem and progenitor cell transplantation**

Human HSCs were injected intravenously into female FNRG or FNRG/A2 xenorecipient mice 10-14 days after human hepatocyte transplantation, to achieve dual engraftment, or into non-hepatocyte-transplanted FNRG or FNRG/A2 mice. Xenorecipients (8-12 weeks of age at the time of HSC injection) were preconditioned through irradiation with 300 cGy 4-6 h before HSC injection. Male and female mice, transplanted with CD34+ HSCs derived from various human donors, were used in this study.

**HBV DNA and HIV RNA isolation from supernatants**

HBV DNA was isolated using the QIAamp MinElute Virus Spin Kit (Qiagen, Hilden, Germany). HIV-1 RNA was purified from plasma with the QIAamp Viral RNA Mini Kit (Qiagen, Hilden, Germany).

**Total HBV DNA isolation from liver tissue**

Liver tissue from HBV-infected FNRG/A2 mice was preserved in RNAlater (Thermo Fisher Scientific, Waltham MA) once excised, and stored at -80^0^C. To isolate total HBV DNA, 100 µl lysis buffer (50 mM Tris-Base, 50 mM EDTA, 1% SDS, 100mM NaCl pH 8.0) was added to 20-25 mg tissue The sample was further digested by addition of 20 µl Proteinase K per sample from a QIAamp DNA Mini Kit (Qiagen, Hilden, Germany) for 18 h at 37ºC. 1 µl of RNase A (SigmaAldrich, St. Louis, MO) was then added to the lysate and incubated at room temperature (RT) for 2 min. 500 µl AL lysis buffer (Qiagen, Hilden, Germany) were subsequently added to the lysate, and samples were then incubated at 70^º^C for 4 h, with vortexing every 20 min. 500 µl absolute EtOH were then added and mixed thoroughly with samples by inverting 10 times. This suspension was then applied to a QIAamp DNA Mini Kit column and centrifuged for 1 min at 13,000 rpm. The samples were centrifuged again in new tubes for 1 min at 13,000 rpm. DNA was eluted in 50 µl AE buffer and concentrations measured using a NanoDrop Spectrophotometer (Thermo Fisher Scientific, Waltham, MA).

**Isolation of HBV cccDNA from total HBV DNA**

A 25-µl aliquot of total HBV DNA extracted from each liver specimen was digested with 1 µl plasmid-safe DNase (Epicentre, E3101K, Madison, WI) to degrade all chromosomal DNA, along with any linear HBV DNA. Per the manufacturer’s instructions, the reaction mix was incubated at 37^º^C for 30 min. Following digestion, plasmid-safe DNase was heat-inactivated by incubating samples at 70^º^C for 30 min. The cccDNA was then purified using a DNA Clean-up and Concentration Kit (Zymo, Irvine, CA), eluted in 30 µl sterile ddH_2_O, and the residual amount of DNA quantified using a NanoDrop Spectrophotometer. Samples were either used immediately for HBV cccDNA quantification by qPCR, or were stored at -20^0^C.

**Total RNA isolation from liver tissue**

Liver tissue was weighed and preserved per the steps taken for DNA extraction from liver tissue. Total RNA was extracted using an RNeasy Kit (Qiagen, Hilden, Germany). Stainless steel beads (5 mm, Qiagen, Hilden, Germany) and 350 µl RLT buffer were added to each Sample Tube RB (2ml, 990381, Qiagen, Hilden Germany). Samples were then homogenized using a tissue disruptor (Fisher Scientific, Hampton, NH). Samples were eluted twice, first by the addition of 30 µl RNase-free water to columns and centrifugation of the samples for 1 min at 10,000 rpm, followed by another addition of 50 µl RNase-free water and centrifugation, as before.

**Generation of HBV stocks**

HepG2.2.15 cells ^5^ were cultured in DMEM F12 media supplemented with 10% FBS, 1% Pen-Strep. Media from the HepG2.2.15 culture was collected every two to three days for approximately 3 weeks. The collected media was sterile-filtered through a 0.22-μm filter (Millipore, Darmstadt, Germany) and was then concentrated 100-fold using a stir cell concentrator (Millipore, Darmstadt, Germany). After concentration, the virus was processed with a HiTrap heparin column (GE, Fairfield, CA) to further concentrate and purify infectious virus particles from non-infectious sub-viral particles. The column was washed with 5 column volumes of wash buffer (20 mM phosphate buffer, 50 mM NaCl, pH= ~7), and the virus was eluted with elution buffer (20 mM phosphate buffer, 2 M NaCl, pH= ~7). The viral stock was then dialyzed using a dialysis cassette (Millipore, Darmstadt, Germany). After dialysis, virus was aliquoted into cryovial tubes and cryopreserved at -80°C until further use. The virus was passaged through a human liver chimeric mouse, resulting in high viremia. Serum was collected and diluted 1:20. A 200-µl aliquot of this 1:20 diluted virus was then used to infect all FNRG/A2 mice. The same viral stock was used for all experiments.

**Generation of HIV stocks**

The CCR5-tropic strain of HIV-1 (JR-CSF) was generated by transfection of 293T cells (ATCC) with plasmid containing full-length HIV-1 (JR-CSF) genome as previously described ^6^.

**Expansion of human myeloid cell lineages and NK cells in FNRGF/A2 mice**

Prior to viral infection, highly engrafted FNRGF/A2 mice (serum albumin concentration ≥1 mg/ml; hCD45+ cells ≥50%) will be injected intravenously with a recombinant adenovirus expressing human Flt3LG (AdV-Flt3LG) (5 x 10^10^ GE / animal)

**Infection of mice with HBV and HIV**

Mice were injected intraperitoneally (i.p) with 1 x 10^7^ GE/animal HBV genotype D subtype ayw (200-µl volume) from HepG2.2.15 cells that were passed once through a human liver chimeric FNRG mouse. FNRGF/A2 mice were infected with HBV gt-D between 5-7 days post-AdV-Flt3LG administration. For HIV infection, mice were anaesthetized and infected with HIV-1 (JR-CSF) (10 ng p24 per mouse) via retroorbital injection.

**Detection of HBV DNA by qPCR**

The following primers and probes were used for amplification of HBV DNA: 5’-CCGTCTGTGCCTTCTCATCTG-3’ (forward primer), 5’-AGTCCAAGAGTCCTCTTATGTAAGACCTT-3’ (reverse primer), and 5’-FAM-CCGTGTGCACTTCGCTTCACCTCTGC-TAMRA-3’ (probe) (IDT, San Jose, CA). The probe and primers were used at a final concentration of 300 nM and 600 nM, respectively. A master mix, made with 2X TaqMan reaction mix (AppliedBiosystems, Foster City, CA), primer/probe mix, and ddH_2_O, was added to wells of a 96-well plate. 5 µl of each sample (isolated from either mouse serum or liver DNA) and DNA standards were added to their respective wells. The following PCR program was run on a StepOnePlus Real-Time PCR System (Life Technologies, Carlsbad CA): 50^º^C for 5 min, 95^º^C for 10 min, followed by 40 cycles of 95^º^C for 15 sec, 56^º^C for 40 sec, and 72^º^C for 20 sec. Lastly, a melt curve was performed at 95ºC for 10 sec, 65^º^C for 10 sec, 50^º^C for 5 sec, and 95^º^C for 5 sec.

**Detection of HBV pre-genomic RNA by qPCR**

A modified iTaq Universal SYBR Green One-Step qPCR kit (BioRad, Hercules, CA) protocol was used to quantify HBV pgRNA. A primer mix with each primer at 3 µM was created with the forward primer 5’-GAGTGTGGATTCGCACTCC-3’ and the reverse primer 5’-GAGGCGAGGGAGTTCTTCT-3’ (IDT, San Jose, CA). A master mix was created per reaction as follows: 5 µl SYBR mix, 0.125 µl RT, 1 µl primer mix, and 1.875 µl ddH_2_O. The following cycling conditions were used: 50ºC for 10 min and 95^º^C for 1 min, followed by 40 cycles of 95^º^C for 15 sec, and 60^º^C for 1 min. The melt curve was performed at 95^º^C for 5 sec, 65^º^C for 5 sec, 95^º^C for 15 sec, and 50^º^C for 5 sec.

**Detection of HBV cccDNA by qPCR**

The following primers and probes were used for amplification of HBV cccDNA: 5’-GTCTGTGCCTTCTCATCTGC-3’ (forward primer), 5’-AGTAACTCCACAGTAGCTCCAAATT-3’ (reverse primer), and 5’-FAM-TTCAAGCCTCCAAGCTGTGCCTTGGGTGGC-TAMRA-3’ (probe) (IDT, San Jose, CA). The final concentrations of DMSO, the probe, and primers were 4%, 0.2 µM, and 0.9 µM, respectively. A 5-µl aliquot of HBV DNA, isolated from either mouse serum or liver, was used per replicate. The following qPCR protocol was used: 95^º^C for 10 min, followed by 50 cycles of 95^º^C for 15 sec, and 61^º^C for 1 min. To confirm that HBV cccDNA specifically was indeed being amplified, primers that are biased for HBV cccDNA amplification over rcDNA were used: 5’-GCCTATTGATTGGAAAGTATGT-3’ (forward primer), 5’-AGCTGAGGCGGTATCTA-3’ (reverse primer) (IDT, San Jose, CA) resulting in a 1,100-bp amplicon^7^. The following PCR cycle was used: 98^º^C 30 sec, followed by 36 cycles of 98^º^C for 10 sec, 61^º^C for 30 sec, and 72^º^C for 45 sec, followed by a final extension step of 72^º^C for 2 min.

**Detection of HIV RNA by RT-qPCR**

HIV RNA was reverse transcribed and quantitatively detected by real-time PCR using the TaqMan Fast Virus 1-Step PCR kit (ThermoFisher Scientific). The primers used for detecting the HIV Gag gene were: 5′-GGTGCGAGAGCGTCAGTATTA AG-3′ (forward primer) and 5′-AGCTCCCTGCTTGCCCATA-3′ (primer). The probe 5’-FAM-AAAATTCGGTTAAGGCCAGGGGGAAAGAA-QSY7-3’ used for detection was ordered from Applied Biosystems. Reactions were set up per the manufacturer’s guidelines and were run on the QuantStudio 6 Flex PCR system (Applied Biosystems).

**mRNA expression analysis**

0.2-2 µg of total RNAs were used for cDNA preparation by reverse transcription with random primers and SuperScript III (Invitrogen), per the manufacturer’s instructions. 2 µl of diluted (1:10) cDNA were used for quantification with the Power SyBR Green PCR Master Mix (Applied Biosystems) on the QuantStudio 6 flex (Applied Biosystems, Foster City, CA), using specific primers. Data were normalized using human GAPDH as the housekeeping gene and expressed as the relative mRNA level compared to the controls. Primer sequences (forward; reverse) were as follows:

TGF-β: 5’-GACATCAACGGGTTCACTACCG-3’; 5’-AGAAGCAGGAAAGGCCGGTT-3’

GAPDH: 5’-GGAGTCAACGGATTTGGT-3’;5’-AAGATGGTGATGGGATTTCCA-3’

CD163: 5’-GGGCTAATTCCAGTGCAGGT-3’; 5’-GCTGACTCATTCCCACGACA-3’

ISG15: 5’-CGCAGATCACCCAGAAGATCG-3’; 5’-TTCGTCGCATTTGTCCACCA-3’

IFITM3: 5’-ATGTCGTCTGGTCCCTGTTC-3’; 5’-GTCATGAGGATGCCCAGAAT-3’

Mx-1: 5’-GGTGGTCCCCAGTAATGTGG-3’; 5’-CGTCAAGATTCCGATGGTCCT-3’

IFN-β: 5’-GTGCCTGGACCATAGTCAGAGTGG-3’; 5’-TGTCCAGTCCCAGAGGCACAGG-3’

ARG1:	5’-GTGGACAGACTAGGAATTGGC-3’; 5’-TCCAGTCCGTCAACATCAAAAC-3’

Timp1:	5’-CTGTGAGGAATGCACAGTGTTT-3’;	5’-TCCGTCCACAAGCAATGAGT-3’

HBV pgRNA:	5’-GAGTGTGGATTCGCACTCCTC-3’; 5’-CGAGGCGAGGGAGTTCTTCT-3’

**HBsAg ELISA**

Detection and quantification of HBsAg was performed via ELISA, per the manufacturer’s instructions (GS HBsAg EIA 3.1, Bio-Rad, Hercules, CA). Briefly, a 100-µl sample of a 1:20 dilution of supernatant was prepared in 1x DPBS. Absorbance was read at 450λ on a BertholdTech TriStar (Bad Wildbad, Germany).

**Assessment of human hepatocyte engraftment by human albumin ELISA**

Human albumin in mouse serum were quantified by ELISA. 96-well flat-bottomed plates (Nunc, Thermo Fisher Scientific, Walttham, MA) were coated with goat anti-human albumin antibody (1:500, Bethel) in coating buffer (1.59g Na_2_CO_3_, 2.93g NaHCO_3_, 1L dH_2_O, pH = 9.6) for 1 h at 37˚C. Plates were washed four times with wash buffer (0.05% Tween 20 (Sigma Aldrich, St. Louis, MO) in 1x DPBS), and then incubated with SuperBlock Blocking Buffer (Fisher Scientific, Hampton, NH) for 1 h at 37˚C. Plates were then washed twice. Human serum albumin (Sigma Aldrich, St. Louis, MO) was diluted to 1 µg/ml in diluent (10% SuperBlock Blocking Buffer, 90% wash buffer), then serially diluted 1:2 to establish an albumin standard. Mouse serum was serially diluted 1:10. Coated plates were incubated for 1 h at 37˚C, then washed three times. 50 µl mouse anti-human albumin antibody(Abcam, Cambridge, UK), diluted 1:2000, was added and plates were incubated for 2 h at 37˚C. Plates were washed four times and 50 µl goat anti-mouse-HRP (LifeTechnologies, Carlsbad, CA), diluted 1:10,000, were added and incubated for 1 h at 37˚C. Plates were washed six times. 100 µl TMB substrate (Sigma Aldrich, St. Louis, MO) were added, and the reaction was stopped with 12.5µl 2-N H_2_SO_4_. Absorbance was read at 450λ on a BertholdTech TriStar (Bad Wildbad, Germany).

**H&E staining**

4-µm specimens were sectioned from formalin-fixed, paraffin-embedded (FFPE) tissues, and H&E staining was performed with a Leica ST5020 Slide Stainer. Paraffin was removed using xylene, followed by graded EtOH dehydration. Sections were rehydrated with tap water and stained with Hematoxylin II for 2 min. Slides were clarified for 15 sec with tap water and blued for 2 min. Sections were washed in tap water and, after an EtOH rinse, stained with Eosin Y for 1.5 min. Sections were then dehydrated in graded EtOH, cleared with xylene, and mounted. All staining components were purchased from Thermo Fisher Scientific, Waltham, MA.

**FAH staining**

Immunohistochemistry was performed on FFPE human chimeric murine liver tissues using mouse anti-human (clone 2) FAH (Abcam, Cambridge, UK). Briefly, sections were deparaffinized in xylene (three changes), rehydrated (three changes 100% EtOH, three changes 95% EtOH), and rinsed in distilled water. Antibody incubation and detection were carried out on a NEXes instrument (Ventana Medical Systems, Oro Valley, AZ) using Ventana’s reagent buffer and iVIEW Detection Kit unless otherwise noted. Endogenous peroxidase activity was blocked with hydrogen peroxide. Heat-induced epitope retrieval was performed in a 1200-W microwave oven at 100% power in 10-mM sodium citrate buffer (pH 6.0) for 20 min. Sections were allowed to cool for 30 min and then rinsed in distilled water. Mouse antihuman FAH was diluted 1:800 in Dulbecco’s Phosphate-Buffered Saline (DPBS) (Invitrogen, Life Technologies, Carlsbad, CA), and incubated overnight at RT. Primary antibody staining was performed with biotinylated goat anti-mouse antibody, followed by application of a streptavidin-HRP conjugate. The complex was visualized with 3,3-diaminobenzidine and enhanced with copper sulfate. Matched Ig isotype, at equivalent concentration and diluted in DPBS, was used as a negative control. After staining, slides were washed in distilled water, counterstained with hematoxylin, and dehydrated and mounted with permanent medium. Stained slides were scanned at 40X magnification using a Leica Microsystems SCN 400F Whole Slide Scanner. Images were viewed and captured using a SlidePath Digital Image Hub (Leica Microsystems, Wetzlar, Germany).

**Isolation of immune cells from spleen tissue**

Mice were anaesthetized by intraperitoneal injection of a mixture of 100 mg kg^-1^ ketamine and 10 mg kg^-1^ xylazine. Spleens were excised and placed in cold, sterile 1X DPBS for transport. Lymphocytes were isolated from the spleen by cutting the spleen into small pieces using a razor blade and digesting with 5 ml 0.1% collagenase (Sigma-Aldrich, St. Louis, MO) for 30 min at 37**°**C, with vigorous pipetting of samples at 15 min to separate cell aggregates. The cell suspension was then passed over a 100-μM cell strainer to remove any large cell aggregates, and then centrifuged at 1,200 rpm for 5 min at 4**°**C. Cell pellets was then resuspended in 2 ml lysis buffer (BD, Franklin Lakes, NJ), and samples were incubated for 15 min in the dark at RT. The lysis reaction was quenched by the addition of complete DMEM (DMEM, supplemented with 10% FBS, 1% Pen-Strep). Samples were then centrifuged at 1,200 rpm for 5 min at 4**°**C. Samples were washed twice with 10 ml sterile 1X DPBS, and resuspended in 1X DPBS1 x 10^6^ splenocytes were added to each well. Live/dead staining was carried out usingZombie UV Viability Dye (Biolegend, San Diego, CA) for 20 min at RT in the dark. The reaction was quenched by addition of 150µl FACS buffer (1X DPBS, 1% FBS). Splenocytes were then stained with antibody panels, as described below.

**Isolation of immune cells from whole blood**

Mice were bled by cardiac puncture. Blood was collected in EDTA tubes (Sarstedt, Nümbrecht, Germany). Whole blood was centrifuged at 3,500 rpm, for 10 min at RT. Serum was then collected and stored at -80°C until further use. Pellets were lysed in 2 m erythrocyte lysis buffer in the dark for 15 min at RT. Lysis was stopped by addition of 2 ml complete DMEM. Samples were then centrifuged for 5 min at 1,300 rpm at 4 °C. Supernatants were discarded, and cells washed twice with 2 ml sterile FACS buffer. Cells were resuspended in 300 µl FACS buffer and 100 µl per sample were aliquoted into separate wells of a 96 well plate. Plates were centrifuged at 1,500 rpm for 5 min at 4 °C, and the supernatant was discarded. Samples were then stained with antibodies (Panel 1) or antibodies + HBV HBcAg tetramer (Panels 2 & 3) (see below for details about staining of lymphocytes).

**Isolation of immune cells from liver tissue**

Mice were anaesthetized, as above, and livers were perfused using a butterfly needle and 10-ml syringe with 5 ml sterile 1X DPBS. Livers were excised and placed in cold sterile 1X DPBS on ice for transport. Liver were cut with a sterile razor blade into small pieces in a 10-cm culture dish, and resuspended in 10 ml digestion buffer (0.1% w/v Collagenase Type II (Worthington Biochemical Corporation, Lakewood, NJ), 40-mM HEPES (Sigma-Aldrich, St. Louis, MO), 2-mM CaCl_2_ (Sigma-Aldrich, St. Louis, MO), 2-U/ml DNase I (Thermo Fisher, Waltham, MA), HBSS +/+ (Thermo Fisher, Waltham, MA)). Samples were then incubated at 37°C for 15 min, mixed to resuspend, and incubated at 37°C for another 15 min. Digestion was quenched by addition of a 1:1 ratio of complete DMEM. The liver was then passed repeatedly through a 70-µm cell strainer into a 50-ml conical tube, until a homogenous, single cell suspension resulted. Samples were then centrifuged at 350 rpm for 5 min at 20°C. Supernatants were collected into new 50-ml conical tubes. Lymphocytes were isolated using Lymphocyte Separation Medium (Thermo Fisher, Waltham, MA). Samples were then centrifuged at 1,300 rpm for 20 min at 20°C, with no brake upon deceleration. 2-3 ml of the interphase was collected using a sterile, filter-tipped 1 ml pipet. The interphase was then placed into a new 15-m conical tube and washed twice with complete DMEM. Cells were counted and cell viability was recorded. 2 x 10^6^ cells from each sample were aliquoted into three separate wells of a 96-well plate. Isolated lymphocytes were then stained (see below).

**Flow cytometry and staining**

Cells were incubated with Human Fc block (BD Biosciences, Franklin Lakes, NJ) for 30 min at 4°C. Cells were washed and then stained with an HBcAg tetramer for 30 min at 4°C. Cells were then washed and stained by incubation with master mix containing fluorochorome-conjugated antibodies:

CD45 (BioLegend, clone H130, cat# 304036, Lot# B222308, San Diego, CA)

mCD45 (Biolgend, clone 30-F11, cat# 103114, lot# B219150, San Diego, CA)

CD3 (Invitrogen, cat# MHCD0318, lot# 1386804B, Carlsbad, CA)

CD4 (BD Pharmingen, cat# 555349, lot# 4216811, San Jose, CA)

CD8 (BD Pharmingen, cat# 551347, lot# 5104748, San Jose, CA)

CD19 (eBiosciences, clone HIB19, cat# 61-0199-42, lot# E19-362-101, San Diego, CA)

CD56 (BioLegend, clone HCD56, cat# 318332, lot# B207590, San Diego, CA)

CD33 (BioLegend, clone WM53, cat# 303404, lot# B195145, San Diego, CA)

HLA-DR (eBiosciences, clone LN3, cat# 48-9956-42, lot# 4293352, San Diego, CA)

CD66 (Novus, clone 610F5, cat# NB10077808AF700, lot# A-2-1-12216AF700, St. Louis, MO)

CD38 (eBiosciences, clone HIT2, cat# 61-0389-42, lot# 4289688, San Diego, CA)

CCR7 (eBsiosciences, clone 3D12, cat# 12-1979-42, lot# E13143-112, San Diego, CA)

CD45RA (BioLegend, clone HI100, cat# 304120, lot# B195192, San Diego, CA)

PD1 (eBioSciences, clone eBioJ105, cat# 47-2799-42, lot# 4272759, San Diego, CA)

CD127 (eBiosciences, clone eBRORDR5, cat# 48-1278-42, lot# 4271585, San Diego, CA)

CD28 (eBioSciences, clone CD28.2, cat# 61-0289-42, lot# 4295464, San Diego, CA)

CD27 (eBioSciences, clone 0323, cat# 47-0279-42, lot# 508451-1637, San Diego, CA)

iTAg Tetramer/APC-HLA-A*0201 HBV core FLPSDFFPSV (MBL International, cat# T01032, Woburn, MA)

Flow cytometry data collection was performed using an LSRII Flow Cytometer (BD Biosciences). Data were analyzed using FlowJo software (TreeStar). Human immune cell subsets were gated as follows: T cells, CD45+ CD3+ CD19-; B cells, CD45+ CD3- CD19+; natural killer cells, CD45+ CD3- CD19- CD56+; natural killer T cells, CD45+ CD3+ CD19- CD56+. For characterization of spleen- and liver-resident T-cell activation, human CD3+ CD4+ and CD8+ T cells were analyzed for their expression of HLA-DR, CCR7, CD45RA, PD1, CCR7, CD27, CD28, and CD127. HBcAg specificity of human CD8+ T cells was also assessed via HBcAg FLPSDFFPSV tetramer staining. Fluorophore compensation was performed using an AbC Anti-Mouse Bead Kit (Life Technologies, Invitrogen).

**HBcAg immunohistochemistry staining**

An unconjugated polyclonal rabbit anti-hepatitis B virus core antigen antibody (Cell Marque Corp Cat# 216A-18 Lot# 17099, RRID: AB_1158068, Rocklin, CA), raised against purified lysates of *E. coli* clones containing HBV core DNA, was used for immunohistochemistry ^8-10^. Antibody optimization was performed on murine FFPE, HBV-infected liver tissue, based on known immunohistochemical conditions for HBV-infected human liver samples utilizing the same antibody, and run in parallel. Chromogenic immunohistochemistry was performed on a Ventana Medical Systems Discovery XT instrument (Ventana, Oro Valley, AZ) with online deparaffinization using Ventana’s reagents and detection kits, unless otherwise noted. 4-µm specimens were sectioned from FFPE tissues, and both markers were antigen-retrieved in extended Cell Conditioner 1 (Tris EDTA, 52 min). Samples for surface antigen detection were blocked in DPBS (Invitrogen/Life Technologies, Carlsbad, CA) with 1% non-fat dry milk, 1% BSA, and 0.05% Tween-20 for 30 min. Antibodies against HBcAg were applied neat, and incubated for 2 h at RT. Primary antibody was detected with an anti-rabbit horseradish-peroxidase-conjugated multimer by incubation for 8 min. The complex was visualized with 3,3-diaminobenzidene and enhanced with copper sulfate. Slides were washed in distilled water, counterstained with hematoxylin, and dehydrated and mounted with permanent media. Isotype negative controls, including non-infected murine liver tissue, were run in parallel with experimental samples.

**Statistical analyses**

All statistical analyses were performed with GraphPad Prism 6.0h software. Data are presented as means ± SEM. Multiple group comparisons were analyzed by one-way ANOVA with a Bonferroni’s multiple comparisons test, or by the Kruskal-Wallis test. P values <0.05 were interpreted to be statistically significant.

**REFERENCES**

1.	de Jong YP, Dorner M, Mommersteeg MC, et al. Broadly neutralizing antibodies abrogate established hepatitis C virus infection. *Sci Transl Med.* 2014;6(254):254ra129.

2.	Billerbeck E, Horwitz JA, Labitt RN, et al. Characterization of human antiviral adaptive immune responses during hepatotropic virus infection in HLA-transgenic human immune system mice. *J Immunol.* 2013;191(4):1753-1764.

3.	Waskow C, Liu K, Darrasse-Jèze G, et al. The receptor tyrosine kinase Flt3 is required for dendritic cell development in peripheral lymphoid tissues. *Nat Immunol.* 2008;9(6):676-683.

4.	von Schaewen M, Hrebikova G, Ploss A. Generation of Human Liver Chimeric Mice for the Study of Human Hepatotropic Pathogens. *Methods Mol Biol.* 2016;1438:79-101.

5.	Sells MA, Chen ML, Acs G. Production of hepatitis B virus particles in Hep G2 cells transfected with cloned hepatitis B virus DNA. *Proc Natl Acad Sci U S A.* 1987;84(4):1005-1009.

6.	Ahodantin J, Nio K, Funaki M, et al. Type I interferons and TGF-β cooperate to induce liver fibrosis during HIV-1 infection under antiretroviral therapy. *JCI Insight.* 2022;7(13).

7.	Seeger C, Sohn JA. Targeting Hepatitis B Virus With CRISPR/Cas9. *Molecular therapy Nucleic acids.* 2014;3:e216.

8.	Sharma RR, Dhiman RK, Chawla Y, Vasistha RK. Immunohistochemistry for core and surface antigens in chronic hepatitis. *Tropical gastroenterology : official journal of the Digestive Diseases Foundation.* 2002;23(1):16-19.

9.	Goodman ZD, Langloss JM, Bratthauer GL, Ishak K. Immunohistochemical localization of hepatitis B surface antigen and hepatitis B core antigen in tissue sections. A source of false positive staining. *Am J Clin Pathol.* 1988;89(4):533-537.

10.	Stahl S, MacKay P, Magazin M, Bruce SA, Murray K. Hepatitis B virus core antigen: synthesis in Escherichia coli and application in diagnosis. *Proceedings of the National Academy of Sciences of the United States of America.* 1982;79(5):1606-1610.
